# Variable dose analysis: A novel RNAi-based method for detection of synthetic lethal interactions

**DOI:** 10.1101/176974

**Authors:** Benjamin E. Housden, Zhongchi Li, Colleen Kelley, Yuanli Wang, Yanhui Hu, Alexander J. Valvezan, Brendan D. Manning, Norbert Perrimon

## Abstract

Synthetic sick or synthetic lethal (SS/L) screens are a powerful way to identify candidate drug targets to specifically kill tumor cells but such screens generally suffer from low reproducibility. We found that many SS/L interactions involve essential genes and are therefore detectable within a limited range of knockdown efficiency. Such interactions are often missed by overly efficient RNAi reagents. We therefore developed an assay that measures viability over a range of knockdown efficiency within a cell population. This method, called variable dose analysis (VDA), is highly sensitive to viability phenotypes and reproducibly detects SS/L interactions. We applied the VDA method to search for SS/L interactions with *TSC1* and *TSC2*, the two tumor suppressors underlying tuberous sclerosis complex (TSC) and generated a SS/L network for TSC. Using this network, we identified four FDA-approved drugs that selectively affect viability of TSC deficient cells, representing promising candidates for repurposing to treat TSC-related tumors.

## Introduction

A genetic interaction occurs when the combined disruption of two genes produces a phenotype that differs from that expected based on the effects of each individual gene disruption. One type of genetic interaction in which cell viability is reduced only following combined disruption of two genes but not following disruption of either gene alone is called a synthetic sick or synthetic lethal (SS/L) interaction depending on the severity of the viability effect. SS/L interactions have received considerable interest for the development of drug targets for cancers because targeting of a gene that has a SS/L interaction with a tumor suppressor is expected to specifically reduce viability of tumor cells but leave wild-type cells unaffected (Kaelin 2005; Nijman 2011; Thompson *et al.* 2015). In addition, large-scale knowledge of SS/L interactions can be used to gain functional insight into individual genes and network structures (Tong *et al.* 2001; Boone *et al.* 2007; Costanzo *et al.* 2016).

SS/L screens have been performed covering most of the possible pairwise gene combinations in the yeast *Saccharomyces cerevisiae*, leading to insight into the global molecular wiring of a cell (Costanzo *et al.* 2016). Furthermore, SS/L screens have been performed with the aim of identifying drug targets in cultured mammalian cells, including tumor-derived lines (e.g. Luo *et al.* 2009; Cowley *et al.* 2014; Hart *et al.* 2015; Wang *et al.* 2017). However, SS/L screens in general have suffered from limited reproducibility and have resulted in the identification of relatively few effective drug targets (Kaelin 2005; Cox *et al.* 2014; Downward 2015; Vyse *et al.* 2017). This may in part be due to the noisy nature of high-throughput screens. However, a major contributor to this lack of concordance between studies is likely to be context dependence of cancer cell line dependencies, as illustrated by widely varying responses to existing therapeutic agents (Barretina *et al.* 2012; Garnett *et al.* 2012). Thus SS/L interactions can be classified as ‘hard’ interactions, which function independent of genetic or cellular context and ‘soft’ interactions which may reflect cell context (i.e., the genes expressed in one cell line versus others or growth conditions) (Ashworth *et al.* 2011; Wang *et al.* 2017). Consistent with this, a recent study in which SS/L interaction screens were performed in multiple Ras-dependent and Ras-independent AML cell lines found relatively few dependencies common specifically to Ras-dependent lines, suggesting that the majority of Ras-induced dependencies are specific to an individual context (Wang *et al.* 2017). These issues highlight the need to develop robust SS/L screening methods and assess SS/L interactions across diverse genetic backgrounds in order to find effective drug targets that will function in many contexts.

Towards this goal, we reported previously the use of a cross-species screening approach to identify SS/L interactions with *TSC1* and *TSC2*, the two tumor suppressor genes underlying tuberous sclerosis complex (TSC) (Housden *et al.* 2015, 2017). We performed focused dsRNA screens targeting all kinases and phosphatases in wild-type, *TSC1* and *TSC2* mutant *Drosophila* cells. By comparing between screens, we identified three genes that specifically reduce viability of *TSC1* and *TSC2* mutant cells when knocked down. All three had conserved SS/L interactions with *TSC2* in mouse and human cell lines, illustrating the potential of this approach to identify candidate drug targets relevant to humans. By performing these screens in *Drosophila* and then validating hits in diverse mammalian systems, we hoped to identify core SS/L interactions that were not specific to a single human cell type.

Here, we report an additional advance towards improved SS/L screening methods. Using a similar cross-species screening approach, we performed genome-wide screens to identify SS/L interactions with the *TSC1* and *TSC2* genes. Analysis of screen results and comparison between screens revealed that many SS/L interactions were missed in each screen, consistent with the low rate of reproducibility between previous screens. Further investigation revealed that many SS/L interactions involve essential genes that were likely missed due to overly efficient knockdown, which reduced the viability of both wild-type and TSC1/2 mutant cells. We therefore developed a new RNAi-based screening assay that measures phenotypes over a range of knockdown efficiencies in a single sample. This method improved signal to noise ratio in viability assays by approximately 2.5-fold compared to dsRNA assays and detected 86% of positive control SS/L interactions. Using this method in combination with our previous screen results and other pre-existing datasets, we generated a high-confidence SS/L interaction network surrounding the TSC complex. Finally, using this network we identified four FDA-approved drugs showing selective effects on the viability of TSC deficient cells that may represent promising candidates for drug repurposing to treat TSC tumors. Importantly, all four of these drugs showed conserved effects in three mammalian cell culture models of TSC, including two diverse tumor-derived cell lines, illustrating that this screening approach improves the identification of context-independent candidate therapeutic drugs.

## Results

### SS/L interactions are enriched for essential genes

To identify SS/L interactions with the TSC complex, we performed SS/L interaction screens using two dsRNA libraries. The first targeted 13,099 genes, representing the majority of the *Drosophila* genome. The second library targeted 466 *Drosophila* orthologs of putative targets of FDA-approved drugs. By screening this group of genes with high coverage, we improved the chances of identifying SS/L interactions with genes for which clinically-approved drugs already exist, which therefore may be rapidly repurposed for clinical use to treat TSC tumors. From these two screens, 288 genes were identified that had SS/L interactions with *TSC1* and/or *TSC2* (**Tables S1 and S2**). To confirm the validity of these hits, we selected six genes from the genome-wide (GW) screen with varying confidence levels and tested whether their human orthologs also showed SS/L interactions with *TSC2* in a tumor-derived human cell line. Similar to the high validation rate in mammalian systems observed in our previous studies (Housden *et al.* 2015), all six SS/L interactions were validated in this system (**Figure 1A**).

**Figure 1:**
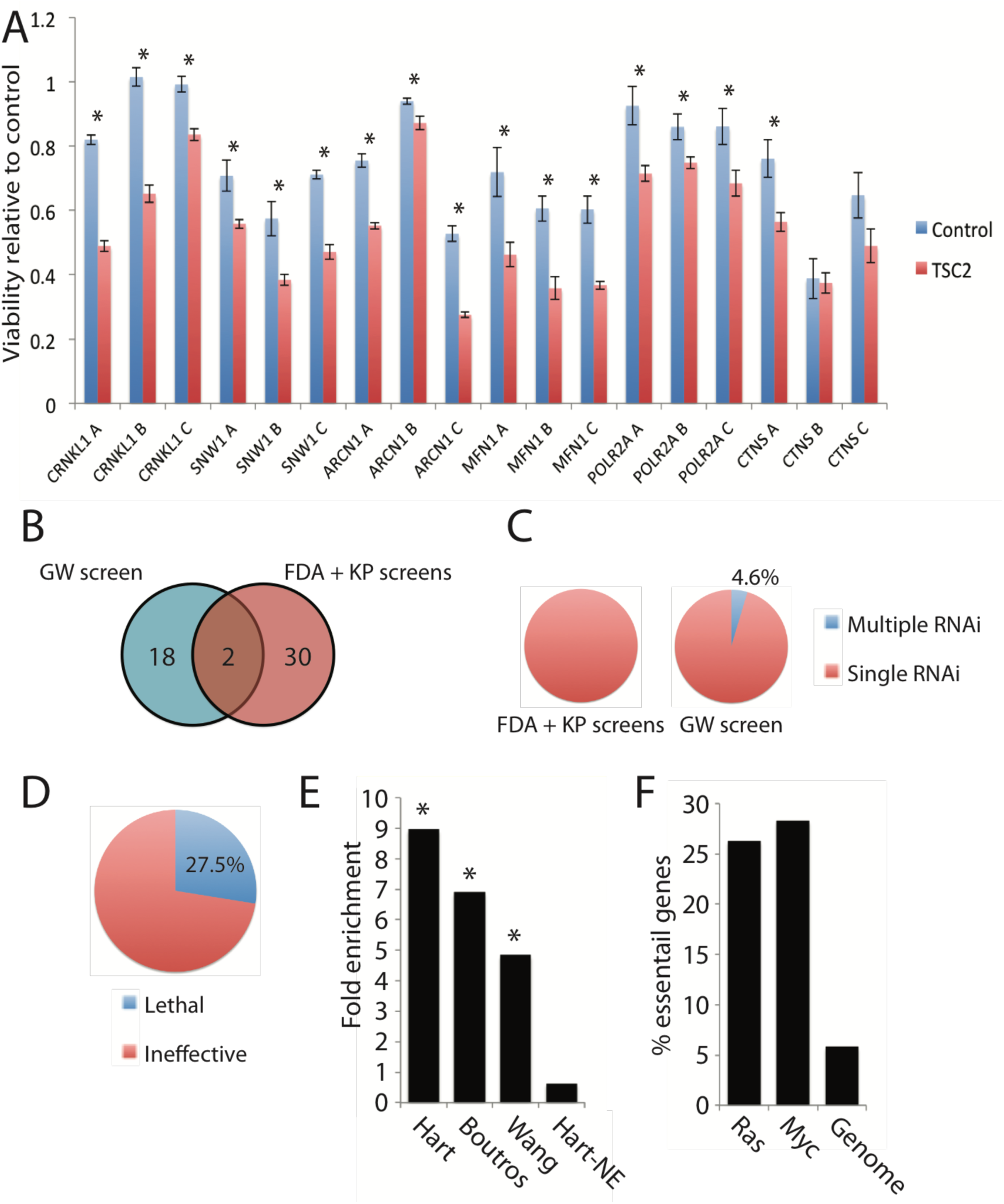
SS/L interactions are enriched with essential genes. **A)** Histogram showing relative viability of *TSC2*-deficient AML-derived cells with either *TSC2* cDNA (blue bars) or empty vector (red bars) addback transfected with the indicated shRNA constructs relative to control shRNA transfection, measured using CellTiter-glo assays. Bars represent average values from nine replicates in each case and error bars indicate standard error of the mean. Asterisks indicate cases where viability of empty vector cells is significantly lower than *TSC2* cDNA addback cells (p<0.05) as determined using t-tests. **B)** Venn diagram illustrating overlap of SS/L genes identified from the genome wide screen (GW screen) and combined FDA and kinase/phosphatase (Housden *et al.* 2015) screens (FDA + KP screens). Only genes that were targeted by both sets of libraries were considered in this analysis. **C)** Pie charts illustrating the proportion of SS/L genes that were identified with multiple independent dsRNA reagents. **D)** Pie chart illustrating screen results for dsRNA reagents targeting SS/L genes that did not detect the SS/L interaction. Reagents were classed as “Ineffective” if no viability effect was detected in wild-type, *TSC1* or *TSC2* cells or “Lethal” if the reagent reduced viability of wild-type cells. **E)** Histogram showing fold enrichment of essential genes amongst SS/L genes identified from the genome wide screen. Three independent datasets were used to define essential (Hart (Hart *et al.* 2014), Boutros (Boutros *et al.* 2004) and Wang (Wang *et al.* 2015)) or non-essential genes (Hart-NE (Hart *et al.* 2014)). Asterisks indicate statistically significant enrichment (p<0.05) based on z-tests to compare with 1000 permutations of randomly selected genes. **F)** Histogram illustrating the % of genes identified as SS/L with Ras or Myc overexpression or activation in previous studies. “Genome” indicates the % essential genes in the whole genome assessed using the same datasets as in E combined to define essential genes (Boutros *et al.* 2004; Hart *et al.* 2014; Wang *et al.* 2015).

Despite the apparent high quality of the SS/L interactions identified, relatively little overlap was observed between the independent libraries screened. For example, a total of 50 SS/L genes were screened in the combined KP/FDA libraries and the genome-wide screen, yet only 2 were identified as SS/L in both datasets (**Figure 1B**). In addition, only 4.6% of the identified SS/L interactions in these screens were reproduced with multiple independent dsRNA reagents despite the fact that 67% of SS/L genes were represented by more than one dsRNA in the screens (**Figure 1C**). This observation is consistent with limited reproducibility in SS/L screens previously performed in mammalian systems. To investigate the reasons underlying the inconsistency between dsRNA reagents and screens, we assessed the effects of dsRNA reagents that did not identify SS/L interactions despite targeting genes that were identified as SS/L using independent dsRNA reagents. We found that approximately 72.5% of reagents had no detectable effect on viability in either wild-type, *TSC1* or *TSC2* mutant cells and likely represent ineffective reagents. By contrast, 27.5% of reagents failed to identify SS/L interactions because they reduced viability of all cell types, suggesting that their targets are essential genes (**Figure 1D and Table S3**). In addition, the 288 SS/L genes identified from the three screens were highly enriched for known essential genes identified in multiple previous studies (**Figure 1E**). These results indicate that genes with SS/L interactions are enriched for essential genes, consistent with previous observations in yeast (Costanzo *et al.* 2016). Furthermore, we found that genes identified from SS/L screens in mammalian systems with activated Ras or Myc, also showed enrichment of essential genes (**Figure 1F**), suggesting that this is a general property of SS/L interactions.

### Variable Dose Analysis (VDA) allows sensitive detection of viability phenotypes

SS/L interactions involving essential genes are detectable within a limited range of target gene knockdown efficiency because weak knockdown is ineffective and strong knockdown is lethal to both wild-type and mutant cells. Therefore, to improve detection of this type of SS/L interaction, an assay is required that allows assessment of viability effects over a range of target gene knockdown efficiencies. To achieve this, we took advantage of the variable efficiency of plasmid transfections between individual cells in a population, resulting in variable plasmid copy number in each cell. By co-transfecting a GFP expressing plasmid and a shRNA expressing plasmid, the GFP intensity can be used as an indirect readout of shRNA expression and therefore target gene knockdown efficiency (**Figure 2A**). We named this method variable dose analysis (VDA). To test the correlation between GFP expression and knockdown efficiency using this approach, we used S2 cells that express mCherry from a genomic transgene insertion into the *CLIC* locus (Neumüller *et al.* 2012). VDA assays were performed in this cell line targeting either a control gene (white) or the mCherry transgene. mCherry fluorescence was then compared to GFP fluorescence and mCherry fluorescence was found to decrease as GFP intensity increased with a non-linear relationship (**Figures 2B and S1**). Therefore, GFP fluorescence is a robust measure of relative target gene knockdown efficiency.

**Figure 2:**
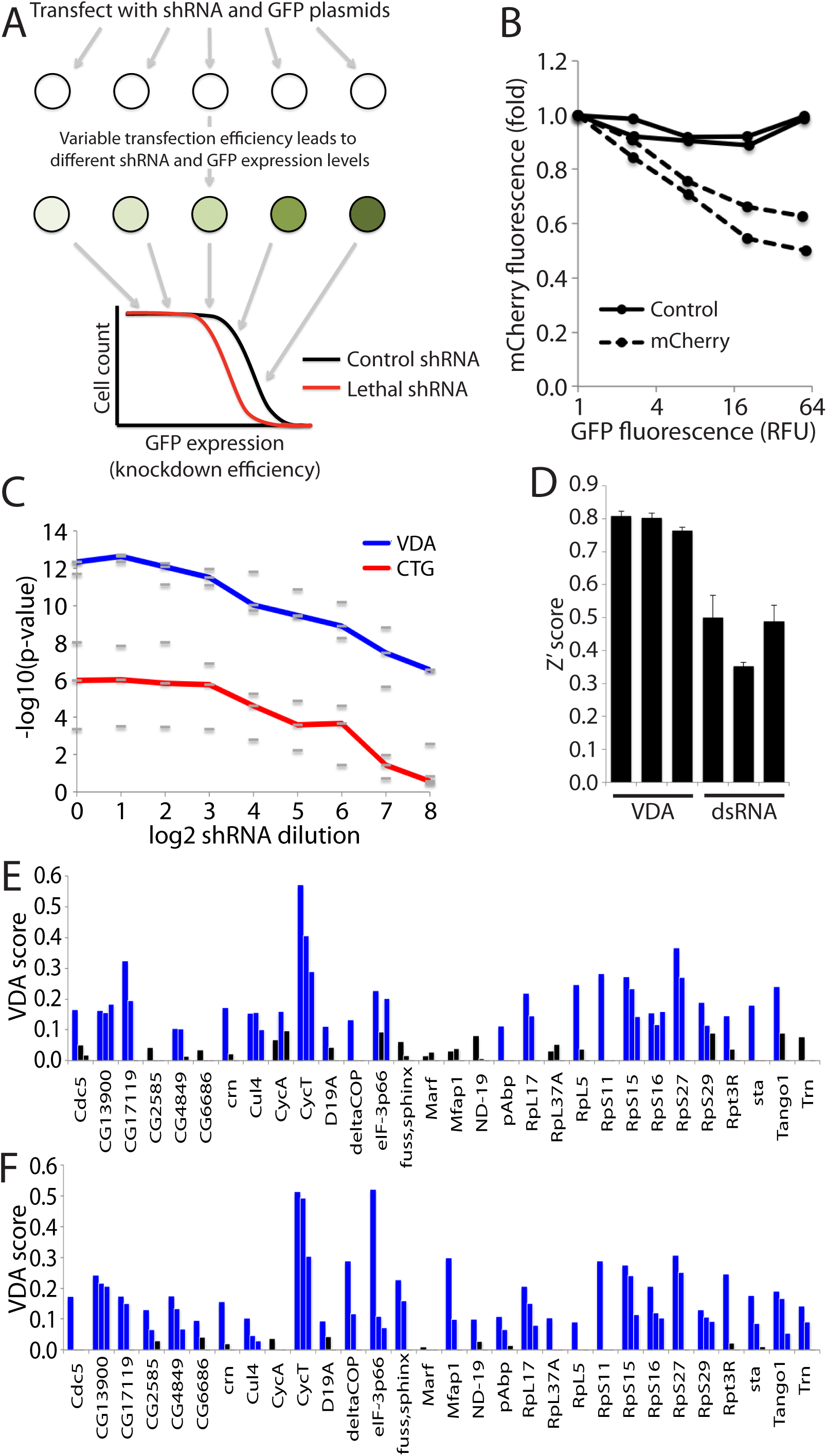
Variable Dose Analysis (VDA) is an effective method to measure viability phenotypes and detect SS/L interactions. **A)** Schematic illustrating the experimental setup of VDA assays. **B)** Graph illustrating the relationship between GFP fluorescence and target gene knockdown efficiency. mCherry fluorescence was used as a measure of knockdown efficiency and is displayed as a fold change to cells with no detectable GFP expression. Each line represents an independent shRNA reagent targeting mCherry (dashed lines) or *white* (solid lines). The graph illustrates results from a single representative experiment from three replicates (see Figure S1 for additional replicates). **C)** Graph illustrating the improved ability of VDA assays (blue line) to detect viability phenotypes compared to CellTiter-glo assays (red line) performed on the same populations of cells. The lines represent median –log10 p-values calculated from six replicate experiments using t-tests. Dashes represent results from three independent groups of six replicates. **D)** Histogram illustrating average Z-prime scores from three independent replicate experiments, each consisting of 30 biological positive control replicates and 30 biological negative control replicate measurements. Each bar represents a different pair of positive and negative control reagents measured using dsRNA/CellTiter-glo or VDA assays as indicated. Error bars indicate standard error of the mean. **E)** Results from VDA assays targeting 30 different genes as indicated. Each bar represents a different shRNA reagent (3 per gene). Blue bars indicate samples with VDA-score greater than 0.1 and black bars represent VDA-scores less than 0.1. **F)** Graph displaying VDA results as in panel E but with VDA analysis performed including cell size correction.

Next, we performed experiments to assess the sensitivity of this method relative to an established viability assay. We co-transfected the GFP reporter plasmid with shRNA plasmids targeting *thread*, an apoptosis inhibitor that robustly induces cell death when inhibited, or a control gene, *white*. Signal strength was varied by serially diluting the *thread* shRNA plasmid with *white* shRNA plasmid. In addition, the same samples were analysed using CellTiter-glo (CTG) assays, a standard readout that has been widely used in high-throughput viability screens. We found that VDA outperformed CTG assays for detection of weak phenotypes and p-values remained highly significant even when the *thread* shRNA was diluted 256-fold compared to the standard dose (**Figure 2C**).

Finally, in order to directly compare VDA with established dsRNA-based methods, we generated three pairs of positive and negative control shRNAs for cell viability targeting *thread* and *white* respectively. We then performed VDA assays in S2R+ cells and calculated Z’ scores for each pair of reagents. In addition, we used three pairs of positive and negative control reagents from the DRSC dsRNA collection (Boutros *et al.* 2004; Hu *et al.* 2017) and performed similar assays using a CTG readout. Comparison between these results showed that the Z’ scores for VDA assays were consistently higher than for dsRNA assays, corresponding to an increase in signal-to-noise ratio of approximately 2.5-fold (**Figure 2D**). In addition, VDA assays had reduced variation between control reagent pairs, indicating that these assays may be more robust to differences in reagent efficiency.

Overall, these experiments demonstrate that VDA is a highly sensitive and robust method for the detection of viability phenotypes.

### VDA assays efficiently identify SS/L interactions with essential and non-essential genes

In order to test whether VDA assays are able to robustly identify known SS/L interactions with both essential and non-essential genes, we generated three shRNA reagents per gene targeting 27 genes identified as SS/L in the dsRNA screens. In addition, to assess the ability of VDA assays to identify SS/L interactions with essential genes, we included 3 genes that were identified as essential (lethal to all cell types) but that had Z-scores at least 1.5-fold lower in both *TSC1* and *TSC2* mutant cells compared to wild-type. Of these 30 genes, 18 were identified as essential genes in previous screens (Boutros *et al.* 2004; Hart *et al.* 2014; Wang *et al.* 2015).

SS/L interactions were identified as shRNA reagents that cause a significantly greater viability reduction in *TSC1* cells compared to wild-type. 70.4% (19/27) of positive control genes were identified as having significant SS/L interactions with *TSC1*, illustrating the high sensitivity of this assay (**Figure 2E, Table S4**). In addition, the three genes that failed to score as SS/L in dsRNA assays due to viability effects in wild-type cells were all identified as SS/L using this assay. Finally, 33% of the genes assessed were identified as SS/L with multiple shRNA reagents (**Figure 2E, Table S4**), indicating a higher rate of reproducibility between reagents compared to dsRNA assays.

Notably, SS/L interactions were not identified for 8 positive control genes. Viability phenotypes were detected for all but one of these genes, indicating that the failure of validation was not due to ineffective reagents. Another possible explanation is that, in addition to affecting cell viability, these genes alter cell size specifically in *TSC1/2* deficient cells and are therefore detected as SS/L in the ATP-based assays used in the dsRNA screens but not in cell count-based VDA assays. To assess this possibility, VDA data were re-analyzed as before but GFP measurements were normalized to cell sizes based on forward-scatter (FSC) readings collected in parallel with GFP fluorescence data. This allows detection of cell size phenotypes as well as viability effects. Using this analysis approach, 28/30 genes tested were identified as SS/L and 19 were identified with multiple reagents (**Figure 2F, Table S4**). This demonstrates the ability of VDA assays to detect multiple different phenotypes in a single assay and to characterize hits in more detail than simple ATP-based assays. Given the improved ability of VDA assays to identify known SS/L interactions compared to dsRNA assays, we used this method to screen 154 genes that can be targeted with well-characterized FDA-approved drugs in wild-type, TSC1 and TSC2 cells. 44 genes were identified as having SS/L relationships with *TSC1* and/or *TSC2* (**Table S5**). Surprisingly, only 1 gene (*porin*) was identified by both VDA and dsRNA assays. TSC1 and TSC2 act together in a protein complex and are thought to share the majority of their functions. Therefore, as SS/L interactions are related to gene function, these genes are expected to have similar SS/L interaction profiles. To assess the relative quality of dsRNA and VDA screens, we therefore compared SS/L interactions identified with *TSC1* or *TSC2* for each method. We selected the top 20 genes from dsRNA and VDA screens in *TSC1* and *TSC2* cells based on either VDA-scores or differences in z-score compared to wildtype cells. For dsRNA assays, 11% (4/36) of top ranked SS/L genes were identified with both *TSC1* and *TSC2*. By contrast, for VDA assays, 33% (10/30) top ranked SS/L genes were shared, suggesting that VDA is a much more robust method for identification of SS/L interactions.

### Integrated analysis of SS/L screen data results in a high-quality SS/L network that is predictive of selective drug effects

Previous studies have shown that genes that physically interact and have related functions share SS/L interaction partners (Tong *et al.* 2001; Costanzo *et al.* 2016). Therefore, in order to identify the most robust hits from the screens and remove false positives, the hits from the dsRNA and VDA screens were pooled (331 genes) and mapped onto high-confidence (score >0.9) protein-protein interaction (PPI) network using the STRING database (Szklarczyk *et al.* 2017) to identify those most likely to have related functions. Due to the relatively low coverage of PPI data, we also manually assigned genes to clusters in cases where gene functions were similar (**Figure 3A, Table S6**). Using this approach, we defined 18 biological processes, each of which were identified as SS/L with the TSC complex by multiple genes. 14 of these processes were supported by both dsRNA and VDA screen results. Furthermore, three of the identified processes have been identified previously as dependencies of TSC deficient cells (e.g. ROS (oxidative stress), proteasome (protein catabolism) and lipid metabolism (Kang *et al.* 2011; Siroky *et al.* 2012; Young *et al.* 2013; Zhang *et al.* 2014; Zhang and Manning 2015; Li *et al.* 2015; Medvetz *et al.* 2015)) indicating that the network is a robust representation of the functional interactions of the TSC complex. Finally, 86.4% (38/44) of the genes identified using VDA assays were assigned to clusters, compared to 68.4% (197/288) of genes identified using dsRNAs, suggesting that VDA is a more robust approach for detection of genuine SS/L interactions.

**Figure 3:**
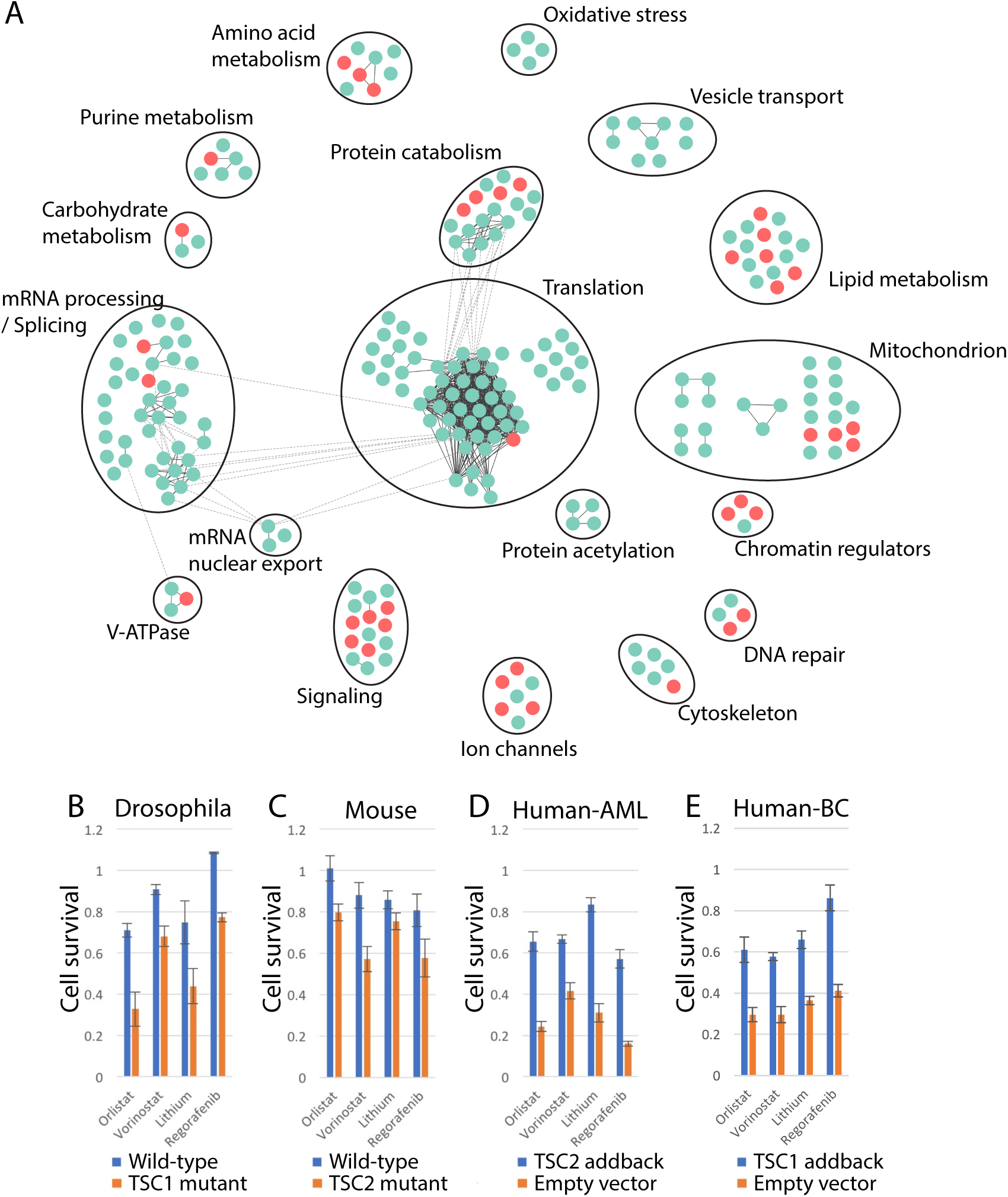
An integrated network analysis approach to identify drugs with selective viability effects on TSC deficient cells. **A)** Network diagram showing genes identified as SS/L with *TSC1* and/or *TSC2*. Circles represent individual genes identified from dsRNA screens (blue) or VDA screens (red). Solid black lines indicate physical interactions within gene clusters and dashed lines indicate physical interactions between components of separate gene clusters. Gene cluster functions were defined based on manual curation of component functions. **B-E)** Histograms illustrating cell viability as measured using CellTiter-glo assays in the presence of the indicated drugs, normalized to vector alone. Bars represent the average of at least six replicate measurements in either wild-type (blue bars) or *TSC1/2*-deficient (orange bars) (B and C) or *TSC1/2*-deficient cells with empty vector (orange bars) or *TSC1/2* cDNA (blue bars) addback (D and E) cells as indicated. Error bars indicate standard error of the mean. Human-AML indicates human AML-derived cells and human-BC indicates human bladder cancer-derived cells.

Genes within the SS/L interaction network represent candidate drug targets to specifically reduce viability of TSC deficient tumor cells. We therefore selected nine FDA-approved drugs that target high-scoring components of the SS/L network and tested for specific viability effects on *TSC2*-deficient *Drosophila* cells. Of these, four had a greater effect on the viability of *TSC2* cells compared to wild-type (**Figure 3B**). In addition, all four of these drugs had conserved effects on *TSC2*-deficient MEFs, *TSC2*-deficient AML derived human cells and *TSC1*-deficient bladder cancer derived human cells (Guo *et al.* 2013) (**Figure 3C-E**), although the quantitative difference in viability effect varied between cell types. Thus, the combined use of improved screening methods with a network-based analysis approach is a powerful method to identify drugs with reproducible viability effects specific to a given genetic background.

## Discussion

SS/L interaction screens have long been considered as a powerful approach for drug target discovery yet have resulted in the identification of relatively few effective drugs. This appears to be due at least in part to a lack of reproducibility between screens despite apparently robust results within studies (Downward 2015). Two technical factors likely contribute to this lack of reproducibility. First, high-throughput screens are inherently noisy, resulting in both false-positive and false-negative results. Second, as we and others have shown (Costanzo *et al.* 2016), SS/L interactions are enriched for essential genes, which are often missed due to overly efficient gene disruption, resulting in general toxicity to all cell types. In particular, this is likely to be an issue for CRISPR screens, which often result in null mutations of the target genes.

In this study, we have addressed both of these issues. First, we developed a novel assay for synthetic lethality called VDA. This approach enables differences in viability between genetic backgrounds to be measured over a range of knockdown efficiencies and can therefore detect SS/L interactions with essential genes at sub-lethal efficiency. In addition, the VDA method is more sensitive and robust to noise than other well-established methods to measure cell viability in *Drosophila* high-throughput screens. Finally, by combining results from two independent screening technologies using a network-based analysis method we have generated a high-confidence SS/L interaction network for the TSC complex. The quality of this network is illustrated by the identification of four effective FDA-approved drugs that represent promising candidates for new therapeutic strategies to treat TSC tumors.

In addition to the technical issues associated with SS/L screens, context dependence of SS/L interactions is also likely to reduce reproducibility between screens in different experimental systems. A common approach to limit this effect is to perform screens in systems that are as closely related to the target tumors as possible, often using tumor-derived cells. More recently, efforts to identify SS/L interactions in panels of divergent cell lines sharing a common tumorigenic driver mutation have been used to identify shared dependencies (Luo *et al.* 2008; Cowley *et al.* 2014; Wang *et al.* 2017). However, this approach requires extensive screening and is limited by high costs and available well-characterized cell lines. Instead, we chose an experimental paradigm in which screens are performed in *Drosophila* cells, which represent a genetic background highly divergent from the target human tumors. Given that many SS/L interactions are highly conserved (Srivas *et al.* 2016), our expectation was that SS/L interactions identified in *Drosophila* cells that could be validated in mouse or human cell lines would represent fundamental context-independent interactions (‘hard’ interactions), which may therefore have a higher success rate when transferred between mammalian systems and to clinical use. Surprisingly, we found that of nine SS/L interactions identified in our *Drosophila* screens that were tested in human cell lines, all could be validated in a highly divergent system (human AML tumor-derived cells) (**Figure 1A** and (Housden *et al.* 2015)). One possible explanation of this is that the increased complexity of mammalian genomes compared to *Drosophila* results in greater network plasticity and therefore more ‘soft’ interactions. In this case, SS/L interactions identified in *Drosophila* have a greater chance of being context independent and therefore more likely to be conserved between divergent systems. This possibility is supported by the similar effects observed for the four identified drugs in diverse backgrounds including mouse and human tumor-derived cell lines. In particular similar effects were observed in bladder cancer-derived cells, which have one of the highest mutation rates of any cancer type (Cancer Genome Atlas Research Network 2014), likely resulting in a highly divergent genetic background compared to the mouse and AML-derived cells.

Previous screening approaches have generally focused on developing the most efficient RNAi reagents possible to maximize resulting phenotypes (Fellmann *et al.* 2013; Kampmann *et al.* 2015; Housden *et al.* 2016). More recently, CRISPR has emerged as a powerful screening technology to generate primarily null mutations in target genes and therefore further increases phenotype strength (Shalem *et al.* 2014; Wang *et al.* 2014). However, these screening paradigms, based on maximizing target gene disruption efficiency, may often be less representative of effects that can be achieved with small molecule inhibitors, which generally incompletely inhibit their targets (Housden *et al.* 2016). It is therefore possible that approaches that are optimized for efficient gene disruption may result in a lower rate of reproducibility with pharmacological assays. In this case, genes identified using VDA may be expected to have more reproducible effects using pharmacological assays because phenotypes can be detected at a relatively low gene disruption efficiency. Consistent with this, three of the four drugs identified in this study were detected only using VDA assays and only one (vorinostat) was identified by both VDA and dsRNA assays. However, neither approach was able to quantitatively predict specific drug effects with no correlation detected between screen score and pharmacological selectivity (**Figure S2**), likely due to differences in the mechanism of disruption between RNAi and small molecule inhibitors (Housden *et al.* 2016). Thus, identification of the most promising targets for pharmacological targeting remains a complex and unresolved issue.

## Materials and Methods

### Construction of the FDA RNAi libraries

We retrieved the FDA drug list from DrugBank (Version 4.0, www.drugbank.ca) and extracted drug gene targets using the DrugBank.xml file. Human drug target genes were mapped to Drosophila genes using DIOPT vs4.0 (Hu *et al.* 2011) and only the high-confident orthologous relationships supported by at least 7 different algorithms were selected for *Drosophila* FDA target libraries. The final FDA library contains 458 *Drosophila* genes. 2 quality amplicons were selected to make the dsRNA library (Hu *et al.* 2017) while 3 different shRNAs were designed based on DSIR tool (Vert *et al.* 2006) and were cloned into Valium20 vector (Perkins *et al.* 2015) to make the shRNA library for VDA assays.

### dsRNA screens

dsRNA screens were performed using the genome-wide, FDA and Kinase/phosphatase libraries available from the *Drosophila* RNAi Screening Center (http://fgr.hms.harvard.edu), following the bathing protocol (http://fgr.hms.harvard.edu/fly-cell-rnai-384-well-format).

### VDA assays

VDA assays were performed by first transfecting a mixture of 10ng actin-GFP, 45ng actin-Gal4 and 45ng shRNA plasmids into wild-type, TSC1 or TSC2 cells seeded into 96-well plates with 30,000 cells per well, following the standard Effectene transfection reagent protocol (Qiagen – 301427). Following 5 days of culture at 25°C, culture plates were analyzed using a BD LSR II flow cytometer. 20,000 events were measured per sample and GFP intensities and FSC measurements were exported for all GFP expressing cells as .csv files for further analysis.

Cytometry data were analyzed using custom Matlab scripts by first normalizing all GFP intensities to cell size measurements (FSC). Next, events were divided into 500 bins based on GFP fluorescence and GFP distributions normalized between all samples. Finally, area under cumulative GFP distribution plots were calculated and compared to negative and positive control samples to calculate viability scores.

Viability scores were calculated as the area between cumulative distribution plots for each sample and median negative control cumulative distribution plot for each plate (based on 5 negative controls targeting the *white* gene per plate) divided by the area between the sample curve and the median positive control curve (based on 5 positive controls targeting the *thread* gene per plate). VDA scores were calculated as viability score in *TSC1* or *TSC2* cells minus viability score in wild-type cells.

The VDA-score cutoff of 0.1 used to identify SS/L interactions was determined by performing 12 replicate VDA assays on each of 6 shRNA reagents determined previously to have no selective effect on TSC deficient cells (Housden *et al.* 2015). We then determined the VDA-score corresponding to a Z-score of 1.5 based on the distribution of VDA-scores from this negative control dataset.

### shRNA assays in human AML tumor-derived cells

Angiomyolipoma cells with stable vector (621-102) or TSC2 (621-103) addback (Li *et al.* 2014) were cultured in DMEM (VWR #45000-312) +10% heat inactivated Fetal Bovine Serum (ThermoFisher Scientific #10437-028) +1X penicillin/streptomycin, (CellGRO 30-002-CI). Cells were transfected with shRNA targeting *ARCN1, CRNKL1, SNW1, CTNS, POLR2A,* or *MFN1* (3 non-overlapping shRNAs per target) in pLKO.1 vector using Lipofectamine 3000 transfection reagent (ThermoFisher Scientific #L3000015) according to manufacturer’s instructions. A scrambled shRNA in pLKO.1 vector was transfected as a control. 6hrs after transfection, cells were washed with 1X PBS and fresh growth media was added. Cell viability was measured 48hr later using the Cell Titer Glo Luminescent Cell Viability Assay (Promega #G7573) according to manufacturer’s instructions. The following shRNA constructs were purchased from the Dana Farber/Harvard Cancer Center Plasmid Information Database (PlasmID): HsSH00157011, HsSH00157016, HsSH00157036, HsSH00167269, HsSH00167262, HsSH00167285, HsSH00152131, HsSH00152139, HsSH00152225, HsSH00116873, HsSH00116854, HsSH00116860, HsSH00129709, HsSH00129751, HsSH00129745, HsSH00157364, HsSH00157314, HsSH00157319.

### Network analysis

SS/L genes identified from dsRNA and VDA screens were pooled and the online STRING database tool (Szklarczyk *et al.* 2017) used to identify high confidence (score ≥ 0.9) protein-protein interactions (PPIs) and cluster the genes. In addition, gene functions were annotated manually based on *Drosophila* or human data in cases where clear orthologs could be identified. Additional SS/L genes were added to network clusters and new clusters created in cases where several genes shared similar functions. The network map was generated using Cytoscape (Shannon *et al.* 2003).

## Acknowledgements

We would like to thank D. Kwiatkowski for the RT4 bladder cancer-derived cell line and E. Henske for the 621-102 and 621-103 AML-derived cell lines. This work was supported by the NIH (P01CA120964), UPenn Orphan disease program (MDBR-15-103-LAM) and the Department of Defense (W81XWH-16-1-0127). N.P. is a Howard Hughes Medical Institute investigator.

